# Transcranial focused ultrasound to rIFG improves response inhibition through modulation of the P300 onset latency

**DOI:** 10.1101/649665

**Authors:** Justin M. Fine, Maria E. Fini, Archana S. Mysore, William J. Tyler, Marco Santello

## Abstract

Response inhibition is important to avoid undesirable behavioral action consequences. Neuroimaging and lesion studies point to a locus of inhibitory control in right inferior frontal gyrus (rIFG). Electrophysiology studies have implicated a downstream event-related potential from rIFG, the fronto-central P300, as a putative neural marker of the success and timing of inhibition over behavioral responses. However, it remains to be established whether rIFG effectively drives inhibition as represented by the P300 activity, and whether rIFG contributions to inhibition are conveyed through either the P300 timing or amplitude. Here, we aimed to causally uncover the connection between rIFG and P300 for inhibition by using transcranial focused ultrasound (tfUS) to target rIFG of human subjects while they performed a Stop-Signal task. By applying tFUS simultaneous with different task events, we found behavioral inhibition was improved only when applied to rIFG simultaneous with a ‘stop’ signal. Applying tFUS simultaneous with the ‘go’ signal or control regions had no impact on behavior. The improvement in inhibition performance caused by tFUS to rIFG during stop conditions occurred through faster stopping times that were paired with significantly shorter P300 latencies, whereas amplitude was not affected. These results reveal a causal connection between rIFG in driving response inhibition in that it may regulate the speed of stopping directly, as indexed by the reduced P300 onset latency during tFUS. Our tFUS-EEG approach provides a causal connection, in healthy humans, between prefrontal rIFG regions and downstream P300 production in service of inhibitory control.

## Introduction

Cognitive control is a process by which a person actively maintains and regulates goal-relevant thoughts and behaviors while suppressing goal or context-irrelevant thoughts and behaviors (Cohen, 2017). The latter, which may entail the stopping of an initiated and/or prepotent action, is referred to as response inhibition (Logan & Cowan, 1984). Response inhibition has been extensively studied as it is central to human behavioral interactions with the environment to prevent adversities and maintain focus on goal-relevant information and responses (Verbruggen & Logan, 2008). Impaired inhibitory control has been associated with several neuropsychiatric disorders, e.g., attention-deficit/hyperactivity disorder (ADHD), impulse control disorders and addiction (Richardson, 2008). Hence, a deeper understanding of the neural dynamics and mechanisms of inhibitory control will contribute to developing better interventional techniques.

To identify neurophysiological markers of response inhibition, many studies have utilized electroencephalography (EEG) and functional magnetic resonance imaging (fMRI). EEG studies have provided substantial evidence about the fronto-central P300 event-related potential (ERP) as a key marker of response inhibition as it tracks the success of inhibitory outcomes (Bekker, Kenemans, Hoeksma, Taslma & Verbaten, 2005; Greenhouse & Wessel, 2013; Kok et al., 2004). Specifically, increased P300 amplitudes have been observed for trials that require response inhibition compared to trials that do not (Enriquez-Geppert et al, 2010). However, the P300 peaks after stop responses, possibly indicating that this amplitude modulation occurs too late to be an indicator of an inhibition response (Huster, Messel, Thunberg & Raud, 2020). Thus, an alternative interpretation is that the P300 amplitude may reflect outcome monitoring (Huster, Enriquez-geppert, et al., 2013) or an attentional orienting process (Corbetta, Patel & Shulman, 2008; Polich, 2007) that likely occurs after inhibition. In contrast, recent evidence has pointed to the P300 onset latency as being a clearer candidate marker of response inhibition for two reasons: it tracks inhibition success and stopping speed (Huster et al., 2020; Wessel & Aron, 2015). Additional evidence for the P300 latency indexing stopping, as contrasted with amplitude, relates to the finding that onset latency modulation occurs before the stopping process has elapsed, while amplitude is modulated after the stopping process (Huster et al., 2020).

Although the P300 itself has a fronto-central topography in EEG studies, hinting at generation from medial prefrontal cortex (MPFC), the areas comprising MPFC have only been linked to inhibitory control as acting downstream from the right inferior frontal gyrus (rIFG). A plethora of fMRI and clinical lesions studies have identified the right inferior frontal gyrus (rIFG) as a central node in triggering response inhibition (for review see Aron, 2011). This would suggest that rIFG might be responsible for the modulation of the P300 signal. However, this putative link between rIFG and P300 modulation remains to be determined. The reason for this gap stems from the issue that EEG and fMRI studies are mostly correlational in nature. These limitations have motivated others to use neuromodulation of brain areas in the inhibitory control network to establish their role. For example, offline repetitive transcranial magnetic stimulation (rTMS) to rIFG during a stop signal task significantly disrupted inhibitory control by increasing Stop Signal Reaction Time (SSRT) (Chambers et al., 2006; Chambers et al., 2007). An EEG-rTMS study further reported that offline stimulation to rIFG reduced right frontal beta power and decreased inhibitory performance, thus establishing right frontal beta as a functional marker of inhibitory control (Sundby, Jana & Aron, 2021). These TMS studies establish rIFG as a central node in the inhibitory control network, however the debate surrounding P300 as a reliable marker of inhibitory control and its connection to rIFG remains unresolved. Additionally, when TMS is applied offline, it becomes challenging to identify the cognitive processing steps that are being perturbed in response inhibition.

Based on the above considerations, we propose that to identify whether the amplitude or onset latency of the P300 relates to inhibition and the involvement of rIFG in this network, it is necessary to perturb rIFG and simultaneously measure the P300. Therefore, the present study was designed to determine whether neuromodulation of rIFG can modify P300 amplitude or onset latency while tracking behavioral inhibition. We simultaneously modulated rIFG activity with transcranial focused ultrasound (tFUS) during EEG recordings of subjects performing a Stop-Signal task (Logan & Cowan, 1984). Based on the above-reviewed evidence provided by EEG and TMS studies, we hypothesized that (1) P300 onset latency is a valid neurophysiological marker of inhibitory control and (2) rIFG is causally related to inhibitory control through modulation of the P300 onset latency. Therefore, we predicted P300 onset latency would track the tFUS-induced changes in response inhibition success and SSRT. We report here that tFUS to rIFG improved stopping behavior through shortening the SSRT and shortening P300 onset latency, thereby causally implicating the role of rIFG in inhibitory control and establishing P300 as a neural signature of inhibitory control.

## Methods and Materials

### Participants

Healthy adult human volunteers (*n* = 63) were randomly assigned to one of three experimental groups. The main experimental group received transcranial focused ultrasound (tFUS) stimulation to right inferior frontal gyrus (rIFG) (*n* = 25; 19 males, mean age = 24.1 yrs., SD = 3.2 yrs.). A second group received stimulation to the ipsilateral somatosensory cortex (*n* = 23; 15 males, mean age = 22.4 yrs., SD = 3.3 yrs.) and was used as the cortical site active control group (S1). A third group received sham stimulation near the right temple (*n* = 15; 8 males, mean age = 24.2 yrs., SD = 2.8 yrs.) and was used as control for possible auditory effects of tFUS modulation (sham rIFG). All individuals were right-handed (self reported) and received financial compensation for participation in the study. Before enrollment, each subject was screened for neurological disorders and a history of epilepsy, stroke, or brain injury. A neurologist from Barrow Neurological Institute (Phoenix, AZ) screened all subjects’ T1 MRI and cleared them before study participation. This study was approved by the institutional review board at Arizona State University. All participants provided informed consent and filled out a safety checklist before commencing the study.

### Behavioral Task and Transcranial Focused Ultrasound design

Response inhibition was assessed using the Stop-Signal Task involving both ‘Go’ and ‘Stop’ trials (**Fig. 1**) programmed in Opensesame (Mathôt et al., 2012). Each trial started with a central fixation cross. In every trial, fixations were replaced by a green ‘Go’ circle (3° x 3° visual angle) after an exponentially-distributed time interval (range: 350-650 ms; mean: 500 ms; standard deviation: 50 ms). Subjects were instructed “to press the up key when detecting the Go circle” (**Fig. 1**). In ‘Go’ trials (rows 1-2, **Fig. 1**), the circle vanished either after the subject’s response or 800 ms elapsed. In ‘Stop’ trials (rows 3-5, **Fig. 1**), the Stop signal was a red square which appeared around the green circle. If the subject successfully inhibited their response with respect to the Stop cue within 800 ms, the red square was extinguished, and the trial was considered a successful inhibition. The time required to inhibit a response following the Stop signal is defined as stop signal reaction time (SSRT) (see below). Timing of the Stop cue relative to Go cue, i.e., the stop signal delay (SSD), was presented at one of four fixed, but subject-specific SSDs. The SSDs were chosen by having each subject perform a practice block of 50 Go trials to determine their baseline Go reaction time (RT). After this block, the 4 SSD levels were set to 25, 35, 75 and 95% of the mean Go RT. These SSDs were fixed throughout the experimental session and were presented in a random order across Stop trials. All trials were separated by a 2-s inter-trial interval (±300 ms random jitter).

**Figure 1.**
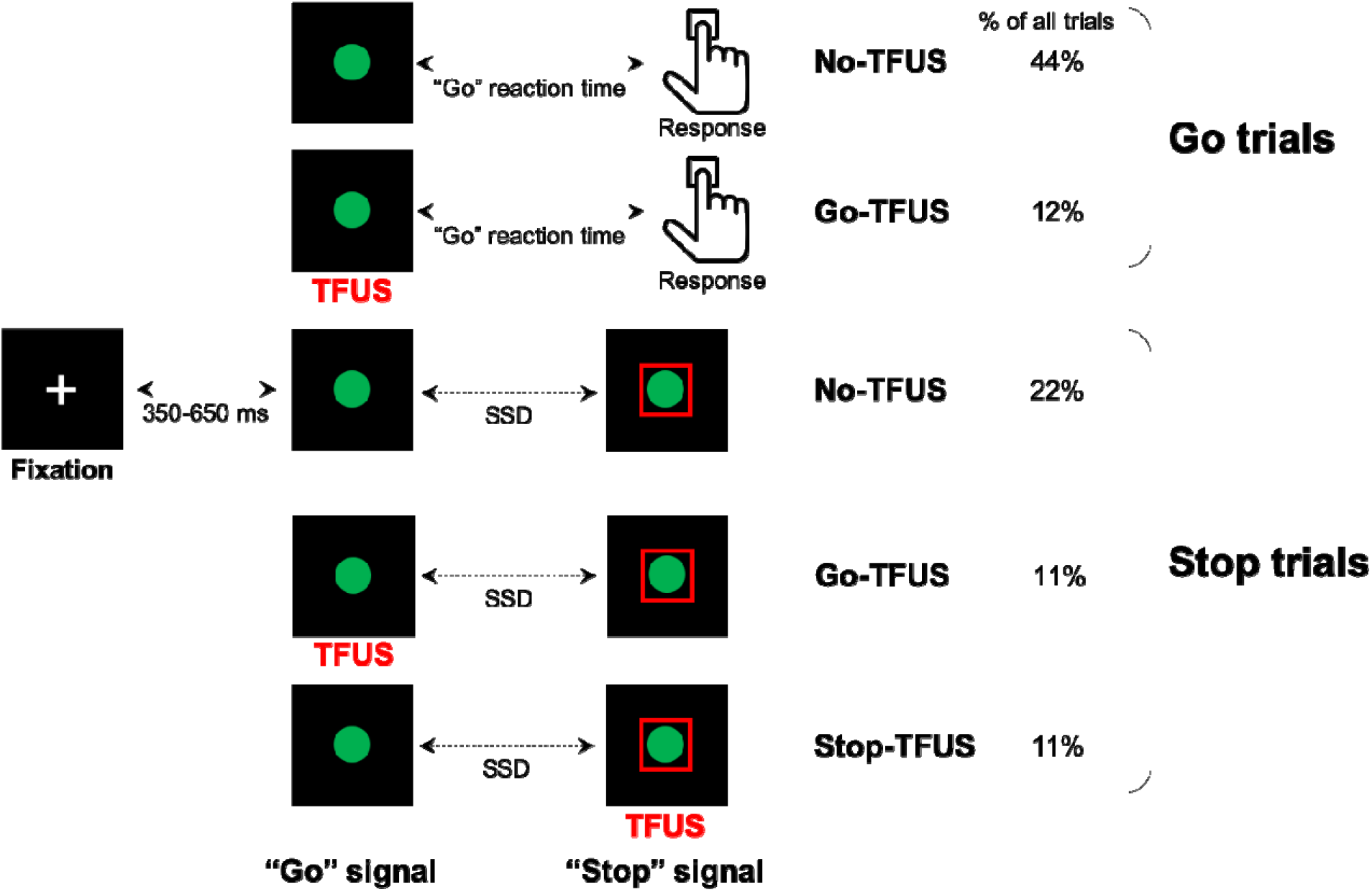
Stop-Signal task and trial types: rIFG group. Each trial type started with a fixation. After a randomly chosen delay (350-650 ms), subjects were asked to respond to a “Go” signal as fast as possible by pressing the up key. On a subset of trials (Stop trials; rows 3-5), a red square appeared at random latencies from the Go signal cueing subject to inhibit their response. Transcranial focused ultrasound (tFUS) was delivered to rIFG for 500 ms either at the onset of the Go (rows 2 and 4) or Stop signal (row 5). SSD: stop signal delay. The same design was used for two control groups (S1 and sham rIFG), although tFUS was not delivered to a cortical site in the sham rIFG group (see text for details). For a subset of Go and Stop trials, no tFUS was delivered (rows 1 and 3).

We delivered Transcranial Focused Ultrasound (tFUS) either simultaneously with the Go signal in both Go and Stop trials, or the Stop signal (**Fig. 1**). This mixture of Go, Stop and tFUS delivery factors generated 5 trial types (**Fig. 1**). The first two consisted of Go trials with no tFUS or with tFUS time-locked to the Go signal (No-tFUS and Go-tFU**S** trials, respectively; rows 1-2, **Fig. 1**). The other three trial types consisted of Stop trials with no tFUS, and tFUS time-locked to either the Go or Stop signal (No-tFUS, Go-tFUS, and Stop-tFUS trials, respectively; rows 3-5, **Fig. 1**). tFUS delivery for Stop trials was evenly distributed across the 4 SSD levels. The overall probability of the occurrence of a Stop trial was 44% of all trials. This proportion of trials accommodates the need for a sufficiently large number of Stop trials to examine tFUS effects on Stop trials across all SSD levels, while enabling a more frequent occurrence of Go than Stop trials (56%; the percentage of each trial type is shown in **Fig. 1**).

Each experimental session consisted of 1200 trials distributed across 12 blocks of 100 trials each. Blocks were segmented into stimulation/no-stimulation and no-stimulation blocks, the former consisting of trials with and without tFUS, and the latter consisting of trials with no tFUS. Trial types (Go and Stop trials) were randomly distributed throughout the experiment. We chose a blocked design to mitigate possible carry-over effects of tFUS across trials. By using two control groups (S1 and sham rIFG), we could determine the extent to which behavioral and/or neural responses associated with tFUS were specific to the target site (rIFG).

### EEG and structural imaging acquisition

#### EEG recording

EEG was recorded using a 64-channel ActiCap system (BrainVision, Morrisville, NC), with 10–20 layout. Data was recorded at a sampling rate of 5 kHz, with 0.1 μV resolution and bandpass filter of 0.1–100 Hz. Impedances were kept < 5 kΩ. Online recordings utilized a ground at AFz and left mastoid reference. At the beginning of each session, electrode layouts with respect to each individual’s head shape were registered using the left and right preauricular, and nasion as fiducial landmarks. This allowed for later co-registration with each individual’s T1 structural MRI scan and for source-localized analysis (see below).

#### Structural MRI (T1)

For guiding tFUS neuronavigation and co-registering EEG electrode placement for source analysis and modeling, we obtained a structural T1 MRI scan for each participant. T1 volumes were collected using an 3D MPRAGE sequence (TR = 2300 ms, TE = 4.5 ms, 1 x 1 x 1.1 mm^3^ voxels, field of view 240 x 256 mm^2^, 180 sagittal slices) in a Philips Ingenia 3T scanner with a 32-channel head coil. Brainsuite was used to process T1s, which included cortical extraction sequence and a surface label-registration procedure with the BCI-DNI atlas. After labeling, we checked the locations and created a mask of either pars opercularis (rIFG group) or the centroid of ipsilateral S1 (S1 group). This volume labeling and mask creation procedure were used for guiding tFUS target identification.

#### tFUS targeting, setup and parameters

A BrainSight neuronavigation system (Rogue industries) along with subjects’ T1 scans were used to guide placement of the focused ultrasound transducer beam profile for stimulation. This was done separately with respect to each individual’s neuroanatomy and mask created from T1s. The first step involved creating a subject-specific mask from cortical atlas registration and projecting into the Montreal Neurologic Institute (MNI) coordinate system. When planning the tFUS target, we considered both MNI coordinates and individual anatomy. For example, metanalysis studies have shown specific activation of the pars opercularis (around x=48, y=16, z=18) for contrasts of successful inhibition versus Go trials and successful versus failed inhibition trials (Chikazoe et al., 2009; Levy and Wagner, 2011). For the rIFG group, we first identified the pars opercularis MNI coordinates. During target planning, we confirmed the coordinates were inside the anatomical region of pars opercularis. We visually confirmed each subject’s pars opercularis tFUS target was rostral to the inferior precentral sulcus and dorsal to the sylvian fissure, and ventral to the inferior frontal sulcus. For the S1 group, tFUS was targeted near MNI coordinates of x=-43, y=-29, z=54 and within the left post-central gyrus.

Before tFUS transducer setup, neuronavigation registered subjects’ T1 scans in virtual space, with their head and the ultrasound transducer in real space. Alignment and cortical registration were performed using nasion, tip of the nose, philtrum, and left and right periauricular notch and tragus as fiducial landmarks. A 3D printed housing held the tFUS transducer, optical trackers, and silicon spacers (ss-6060 Silicon Solutions, Cuyahoga Falls, OH). Acoustic gel was applied to both transducer and scalp. We recorded stimulation target coordinates after placing the transducer in target alignment. In the sham rIFG group, we delivered sham tFUS (Legon et al., 2018) by placing the transducer perpendicular to the rIFG target. For the rIFG and S1 groups, we measured accuracy of stimulation target coordinates by tracking the deviation of the tFUS beam profile from the cortical target throughout the experiment. During the experimental session, we sampled tFUS transducer spatial target deviation during each break. Accuracy was very high, with an average deviation of ±1.5 mm displacement across all subjects and sessions.

Our tFUS setup and parameters were nearly identical to those used in Legon et al. (2014). Briefly, we used a single-element tFUS transducer with a center frequency of 0.5 MHz, focal depth of 30 mm, a lateral spatial resolution of 4.5 mm^2^, and axial spatial resolution of 18 mm^2^ (Blatek). tFUS waveforms were generated using a two-channel, 2 MHz function generator (BK Precision). The system operated by channel 1 produced a pulse repetition frequency (PRF) of 1.0 kHz. Channel 1 also triggered channel 2, which produced short bursts at the 0.5 MHz acoustic frequency. This produced an ultrasound waveform with a carrier frequency of 0.5 MHz, PRF of 1.0 kHz, and duty cycle of 24%. Each stimulation duration was 0.5 s. Transducer power was driven by output from a 40-W linear RF amplifier (E&I 240L; Electronics and Innovation). For the tFUS auditory control in the sham group, we used an auditory recording of the tFUS stimulation clicking sound and delivered it simultaneously with the sham stimulation.

### Computational simulation and validation of tFUS propagation

We quantified peak pressure amplitude, peak intensity and accuracy of the tFUS beam distribution delivered to rIFG using the pseudospectral simulation method in K-wave (Treeby & Cox, 2010). Reference peak pressure planes for the simulations were derived from previous data (Legon et al., 2014). Simulation parameters were first validated by simulating the transducer in water to compare the simulation results with those from previous water tank tests (Legon et al., 2014). The maximum pressure plane at the 30-mm focus was used as a source input pressure for the transducer during the simulation. The transducer was modeled to have a 30-mm radius of curvature. For water simulations, we used a homogenous medium of water density (1000 kg/m^3^) and speed of sound (1482 m/s). We created a computational grid over a 256 x 256 x 256 with 1-mm spacing. The points per wavelength were 6, Courant–Friedrichs–Lewy = 0.1, and simulation time was set to 6 pulses (duration = 250 μs) to ensure simulation stability.

For simulating transcranial ultrasound stimulation, we extracted 3-dimensional maps of the skull from a CT (1-mm resolution) and brain from T1 MRI scans (1-mm resolution) from three preoperative patients at Barrow Neurological institute. The MRI and CT were both co-registered and normalized to the MNI space in SPM12. To mimic our approach of tFUS targeting used in the experiments, we surface registered the gray matter volume to the BCI-DNI atlas and identified the centroid of pars opercularis. The average stimulation location for these three subjects was x = 48, y = 18, and z = 6. This allowed us to map from world coordinates of the scan to MNI coordinates of the target. **Figure 2A** shows T1 and scalp from one subject, together with renderings of the transducer housing, the pars opercularis mask, and the tFUS target applied to all MRIs from the rIFG group. **Figure 2B** shows side views of non-normalized T1s, pars opercularis masks, and the tFUS targets (red dots) for four subjects (**Figure 2B**). Conversion from Hounsfield units in the CT to sound speed and density were done using the relations described in Aubry et al (2003). All skull materials were set using these parameters, while other tissues were treated as homogeneous with parameters set to that of water. Attenuation was modeled as a power law with a β = 0.5 while absorption was modeled with *b* = 1.08 (Treeby and Cox, 2010).

**Figure 2.**
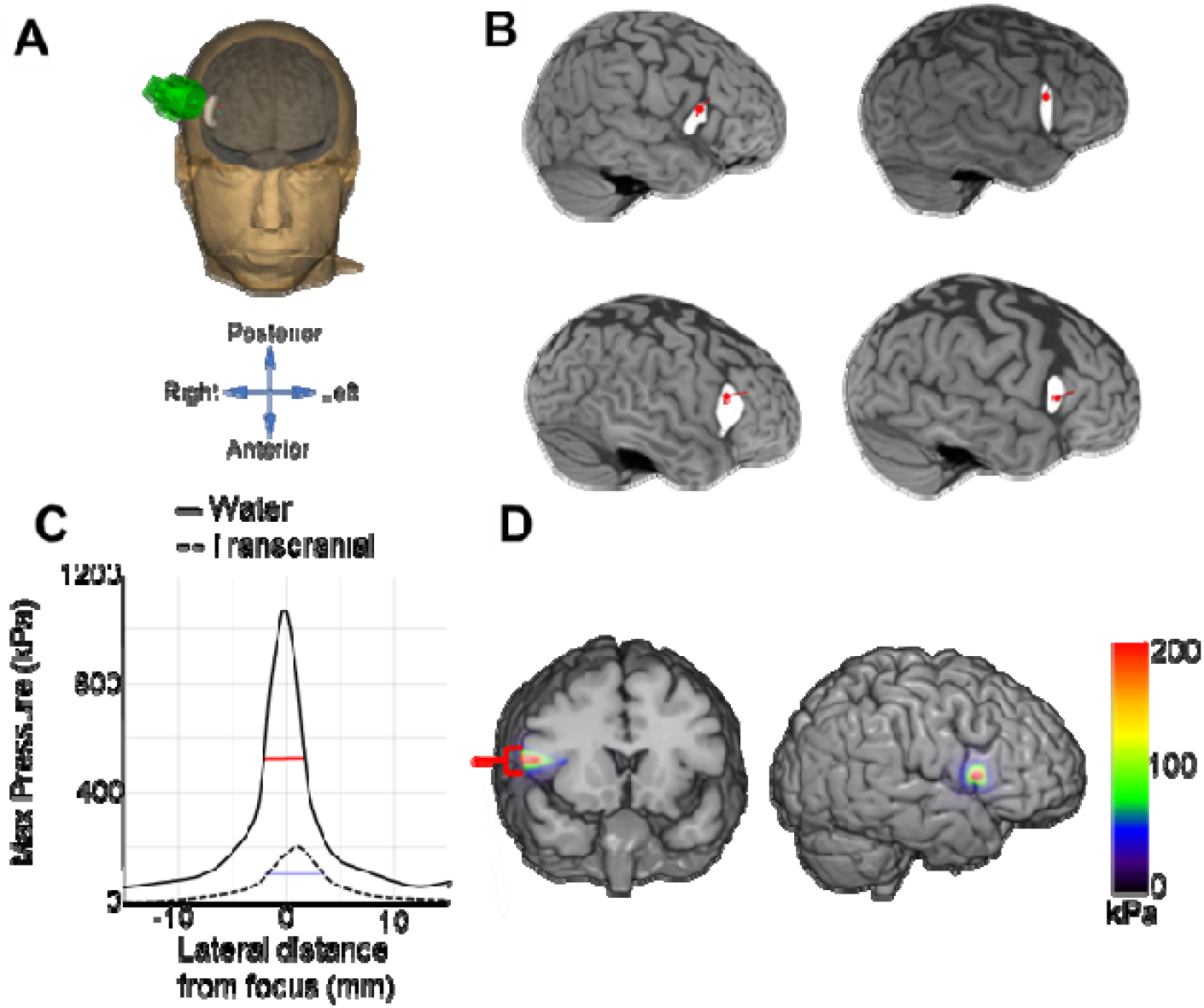
tFUS targeting and simulation. **A**. Average neuronavigation location of tFUS (red dot) applied to all MRIs used in the rIFG group. **B**. Structural brain scans and renderings of the targeted tFUS point (red dot) in pars opercularis (4 subjects). **C**. Lateral maximum pressure profile obtained at 30-mm depth focus in both water and tFUS simulations on a CT scan from one patient (solid and dotted lines, respectively). Horizontal red and blue lines denote full-width half maximum of the spatial profile of lateral pressure. **D**. Simulated transcranial pressure profile onto T1 MRI plot shown as a color overlay.

To assess transcranial stimulation accuracy, the simulated transcranial transmission was compared against simulations of tFUS transmission through water. Differences between these simulations shows the estimated effect of any power absorption and change in acoustic profile after skull transmission. Numerical simulation parameters (see above) were derived to ensure the water simulation here matched the water tank results from a previous study using the same transducer and tFUS experimental parameters (Legon et al., 2014). Simulation of ultrasound through water predicted a max pressure of 1.05 Mpa, spatial peak pulse average intensity (Isppa) of 2 22.4 W/cm^2^ at the focus, and lateral full-width at half maximum of the maximum pressure of 4.39 mm (**Fig. 2C**).

Comparison of simulations and previous water tank data (Legon et al., 2014) using the same transducer and experimental tFUS parameters indicated a 97% match of pressure/intensity at the focus taken over a 5-mm^3^ voxel section in all 3 planes at the focus. Next, modeling of transcranial transmission predicted an average maximum intensity of 2.8 W/cm^2^, which is the intensity range of non-thermal neuromodulation and that exhibits nearly instantaneous measurable effects on EEG (Legon et al., 2014).

Comparing the water and transcranial simulation, accuracy was assessed by comparing shifts in peak pressure. Skull transmission compared to water was shifted 1.25 mm laterally and had a lateral beam profile full-width half maximum of 5.1 mm (**Figure 2C**). These transcranial simulations indicate high spatial precision, with >95% of pressure or energy (kPa) being constrained to pars opercularis in the rIFG group (**Fig. 2D**).

### Statistical analysis

#### Behavioral variables

Our behavioral analyses focused on the following variables: Go trial reaction time (Go RT), percentage of successfully inhibited responses on Stop trials (successful stopping) per SSD, failed inhibition reaction time, and SSRT. The SSRT was estimated using a hierarchical Bayesian parametric approach (Matzke et al., 2013a) that estimates distribution of SSRTs while assuming an ex-gaussian parametric form. We chose this approach as Matzke et al. (2013a) showed that it performs well even when there are only a few trials available per SSD level. This SSRT estimation procedure was run separately per group (rIFG, S1, and Sham rIFG) and trial types (No-tFUS Stop trials, Go-tFUS Stop trials, and Stop-tFUS Stop trials; **Fig. 1**). As we report in the Results section, we used Go RTs combined from Go trials with and without tFUS because stimulation did not alter the Go RT.

The probability of response inhibition changes over levels of SSD, P(respond|signal), and across tFUS conditions were assessed within and across groups by fitting a 2-parameter logistic mixed effects model with random intercepts and slopes to obtain subject- and condition-specific model parameters. P(respond|signal), denoted as *p*, were converted to a negative logit (log((1-*p*)/*p*)) before fitting. As our main goal was to estimate the logistic curve slope (**β**), we ran the mixed-effects model (using LME4 in R) with the full interaction of SSD and stimulation condition in the three Stop trial types (no-tFUS, Go-tFUS, Stop-tFUS). Logistic slopes per subject were estimated by combining fixed and random coefficients. **B** parameters were analyzed using a mixed-design ANOVA on **β** with factors of Group (3 levels) and tFUS (3 levels: No, Go, Stop).

### EEG pre-processing

Continuous EEG data were first down-sampled to 250 Hz, then high-pass filtered (0.1 Hz) and re-referenced to the scalp average. All channels with visually identifiable drift and general noise artifacts for more than 25% of the recording length were removed from further processing. Segments were then removed where channels exhibited muscle and movement activity artifacts (removed portions < 8% of recording lengths across participants). In each of the stimulation groups, the cortical sites of rIFG and S1 were close to the F8 and CP4 electrodes. Therefore, these electrodes could not be used for EEG recording in their groups and were interpolated after artifact removal. Out of all participants, 3 subjects were excluded (2 from S1 group and 1 from rIFG group) from analyses due to EEG recording issues (impedance >25 kΩ across channels). All of these electrodes were then interpolated, and the data were common average re-referenced. To remove other artifactual data segments, we then applied artifact subspace reconstruction for removing or correcting data identified as artifacts; this included outlier noisy segments identified as > 6 standard deviations above the mean. Data were again referenced to ensure a zero-sum voltage across channels.

Independent components analysis was then used to obtain canonical activity maps across electrodes for further analysis, a standard approach in inhibitory control electrophysiology. Each subject’s data were processed with the Adaptive Mixture of ICA (AMICA; Palmer et al, 2008) approach using 1 mixture model. AMICA was chosen because it has shown to have ICA algorithmic superiority in separating data into independent components with dipolar spatial maps (Delorme, Palmer and Makeig, 2014). For each subject, we aimed to select a singular ICA component that has the desired ERPs (N200/P300) that have been putatively linked to inhibitory control (Huster et al., 2014; Huster, 2020; Wessel et al., 2013). To obtain desired Ics, we used a similar approach to previous work analyzing ICA components for electrophysiological markers during inhibition (Wagner et al., 2018; Wessel et al., 2015). Singular equivalent current dipoles were fit for each IC scalp map using DipFit 2.2 in the EEGLAB toolbox. Any IC with a dipole residual variance of less than 15% for predicting the scalp map was removed as these are typically considered to be signals of non-brain activity origin.

Remaining ICs across subjects were then clustered using a K-medoids algorithm. To avoid double dipping in our analysis of ERPs, we only used the features of 3D (X,Y,Z) dipole position and IC weight spatial maps. We optimized selection of the number of clusters using the adjusted rand index (ARI), which measures the stability of the clustering results over multiple runs of the K-medoids algorithm. ARI estimates how similar each cluster is across the runs (from 0 to 1 being identical clusters). We found a maximum ARI of 0.82 for 11 clusters. To select a cluster of Ics for analyzing the N200/P300, we used a previously-employed criterion (Wessel et al., 2016) of requiring it to have: (1) a fronto-radial spatial map topography and (2) a maximal IC weight at 1 of the Fcz, Fz, Cz, Fc1, Fc2 electrodes. For each cluster, some subjects had multiple Ics. In that case, we retained only the subject IC that maximally correlated with the cluster spatial spatial template and had a dipole with a smaller distance from the centroid position from the cluster averaged dipole position.

### GLM statistical model and separating inhibitory from response ERPs

Our analysis focused on a select set of electrophysiological markers that have been linked to the success and speed of stopping. Due to the macroscopic nature of EEG signals, there is likely a temporal overlap of neural processes related to stopping and going during Stop trials. We aimed to separate Go-related activity from Stop-related activity separately for both successfully (SS) and unsuccessfully (US) inhibited Stop trials. This was done by subtracting the Go trial ERPs from the Stop trial ERPs, and using the approach employed by Mattia et al (2012). On a per-subject basis, we found Go-trial ERPs (No-tFUS and tFUS) whose RTs matched those of SS trials based on each subject’s SSRT. These Go trials had to have RTs either equal to or greater than the SSRT. For US stopping trials, we found latency-matched Go trials with RTs by first calculating each subject’s mean signal-respond RT for each of the two highest SSDs. We then calculated the difference in SSD (ms) and searched for Go RT trials for each SSD that fell within the mean signal-respond RT ± half the difference of the SSD (ms). This was done to prevent overlap of activity from both faster and slower Go RTs and signal-respond RTs. These steps were performed separately for the highest and second highest SSD. This procedure was done separately for SS and US trials for both No-tFUS and tFUS conditions. After correcting the SS and US stop trial, the corrected ERPs were averaged across the two highest SSDs per subject (corresponding to 85% and 105% of mean Go RT of each subject). These ERPs were used for the remaining analysis.

### ERP analysis

#### Sensor-space analysis

We examined ERPs at the sensor level using permutation-based dependent samples t-tests that account for spatiotemporal clustering. We performed tests utilizing multiple comparisons with cluster-based p-value corrected at *p* < 0.01, and performed 5000 permutations for each contrast. We used two different contrasts to analyze differences for stopping success overall and tFUS effects on ERPs. The contrasts included comparison of (1) SS –US trials collapsed over No-tFUS and Stop-tFUS conditions, (2) a tFUS effect comparison of SS (No-tFUS) – SS (Stop-tFUS) contrast, and (3) the interaction contrast of SS – US trials with No-tFUS and Stop-tFUS. The first contrast is typically used to determine which areas exhibit ERPs (or brain areas) that differentiate successful inhibition. The second contrast was used to determine which scalp ERPs differentiated successful stopping in the No-tFUS and Stop-tFUS conditions. The third contrast (interaction) was used to determine how the SS-US contrast differed between No-tFUS and Stop-tFUS conditions. The second and third contrasts were the primary focus of our analyses and expected to provide the most direct indication of inhibitory-specific ERP effects. Our reasoning was as follows: if the first SS-US contrast reveals ERP activity and latencies that differentiate successful and failed inhibition, and tFUS alters the potency of inhibitory processes and ERPs that differentiate the level of successful inhibition (second contrast), the interaction contrast should isolate changes in neural response strength to Stop signals and behavioral inhibition outcomes that are jointly modulated by tFUS.

Because recent work has indicated the frontocentral (ERP) P300 onset latency is related to the speed of inhibition (SSRT) across-subjects (Wessel and Aron, 2015), we regressed the between-subject changes in SSRT as a function of the P300 latency change between No-tFUS and Stop-tFUS. To achieve this, we computed the shift in P300 onset crossings between No-tFUS and Stop-tFUS conditions in two steps. First, we took the across-subject mean frontocentral ERP waveform in a time-window of ±50 ms around the zero crossing. To calculate each subject’s zero-crossing time, we calculated the dynamic time warping distance from the template mean ERP to the subject’s ERP. Second, this distance was added to the median zero-crossing time to obtain an individual subject crossing for both the No-tFUS- and Stop-tFUS conditions. These changes in P300 onsets were then regressed against individual subject differences in SSRT between conditions.

## Results

Human participants performed a Stop-Signal task and received online transcranial focused ultrasound (tFUS) on a subset of Go and Stop trials (**Fig. 1**). Subjects were divided into groups according to tFUS stimulation type: (1) an experimental group that received active stimulation to right pars opercularis (rIFG), (2) a control group that received active stimulation to ipsilateral primary somatosensory cortex to account for non-site specific tFUS effects (S1), and (3) a second control group that received sham stimulation to account for possible tFUS-induced auditory artifacts (sham rIFG). Stimulation was applied online at the onset of “Go” or “Stop” signals, separately for both Go and Stop trials (**Fig. 1**). We hypothesized that rIFG is causally related to inhibitory control through modulation of the P300 onset latency. According to this hypothesis, we predicted that the behavioral effects induced by tFUS should be limited to alteration of stopping when tFUS is applied simultaneously with the Stop signal.

### tFUS to rIFG improves stopping behavior

We first addressed how probability of failing to inhibit (P(respond|signal), **Fig. 3A**) changed across tFUS conditions (No-, Go-, and Stop-; rows 3-5, **Fig. 1**) and groups, by fitting a 2-parameter logistic mixed-model to obtain response inhibition curve slopes (**β**) across subjects and tFUS conditions. Analysis of **β** revealed only the rIFG group exhibited a tFUS-altered P(respond|signal), consisting of an enhanced ability to inhibit responses relative to the control groups. ANOVA results indicated a significant Group x tFUS interaction (*F*(4,120) = 3.8, *p* = 0.034, 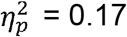). Follow-up one-way within-group ANOVAs across tFUS conditions showed only the rIFG group exhibited differences across the different tFUS conditions (*F*(2,48) = 11.58, *p* = 0.002, 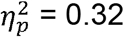). T-tests in the rIFG group showed **β** for Stop-tFUS was lower than No-tFUS and Go-tFUS conditions (both *p* < 0.01: mean **β**’s indicating change in probability for approximately 25% change in normalized SSD: No-tFUS = 0.61 (SE: 0.05; 95% CI: .62-.82), Go-tFUS = 0.65 (SE: 0.09; 95% CI: 0.60-0.82), Stop-tFUS = 0.47 (SE: 0.08; 95% CI: .43-.54)). Therefore, tFUS behavioral effects were limited to the rIFG group and Stop-tFUS trials.

**Figure 3.**
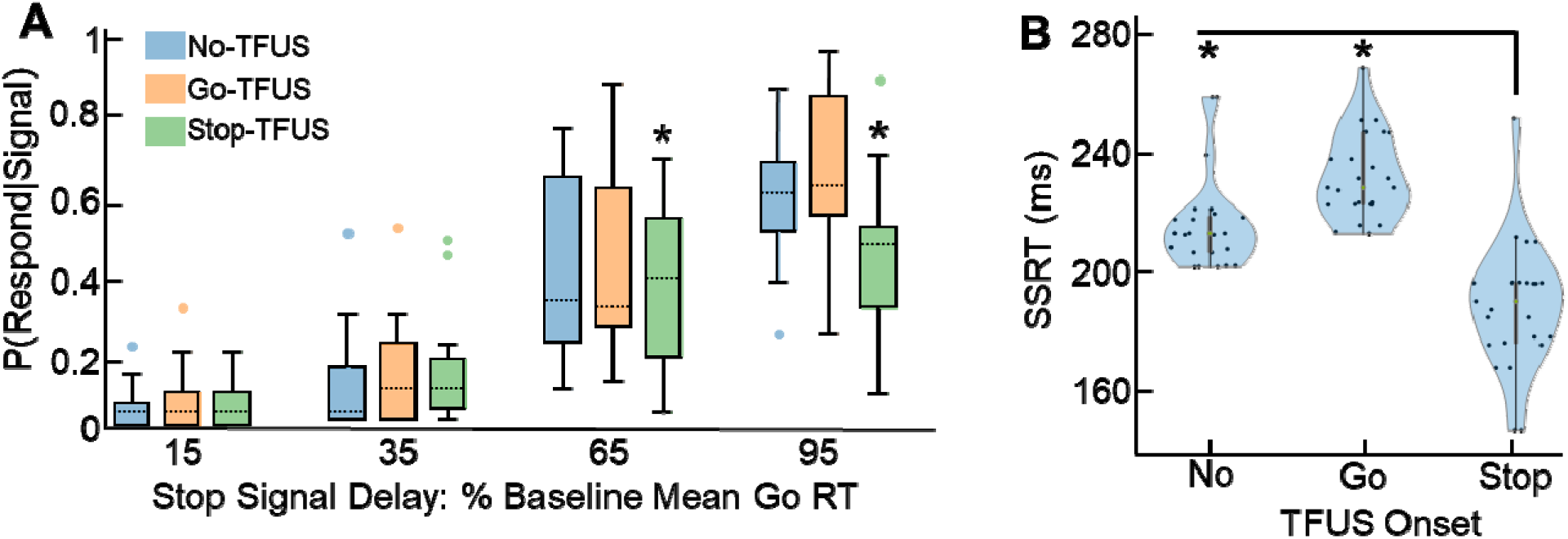
Behavioral effects of tFUS: rIFG group. **A.** Probability of responding across Stop signal delays for tFUS delivered on Stop trials. Dotted lines show median and boxes represent upper and lower quartiles. **B.** Violin plots showing median, interquartile range and kernel density estimated distribution of across-subject distribution of Stop signal reaction times (SSRT) across the tFUS conditions used for Stop Trials.

The effects of rIFG tFUS improvements on successful inhibition performance appear greater at longer SSDs (65% and 95% SSD; **Fig. 3A**). A repeated-measures ANOVA on P(respond|signal) for rIFG group across SSD levels and tFUS onsets supported this by revealing a significant interaction (*F*(6,102) = 8.21, *p* < 0.0001, = 0.33). T-tests between Stop-tFUS and the average of No- and Go-tFUS across all SSDs indicated the interaction resulted from a reduction in P(respond|signal) for Stop-tFUS in the two longest SSDs (both *p* < 0.01; Bonferroni α = 0.0125; mean difference 95% CI: 0.06 – 0.14) These results indicate Stop-tFUS induced improvements of inhibition were more pronounced at longer SSDs (**Fig. 3A**).

Based on the prediction that rIFG is involved in the inhibitory process and our finding that tFUS improved response inhibition only in the rIFG group, we expected tFUS to rIFG-induced changes to P(respond|signal) should result from a shortening of the stopping speed, i.e., SSRT. Notably, tFUS did not affect other behavioral variables, e.g., Go RTs. SSRT analysis in a mixed-design ANOVA indicated a significant Group x tFUS interaction (*F*(4,100) = 10.2, *p* < 0.001, 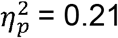). One-way ANOVAs within groups indicated only rIFG group SSRTs (**Fig. 3B**) significantly differed across tFUS conditions (F(2,48) = 23.21, *p* < 0.001), with following t-tests comparing Go-tFUS and Stop-tFUS to No-tFUS confirming that SSRTs were indeed shortest and only altered for Stop-tFUS trials (**Fig. 3B**; mean difference of No-tFUS and Stop-tFUS: 20.24; 95% CI: 10.01 – 30.46). This indicates tFUS to rIFG altered response inhibition by shortening the SSRT.

We used separate analyses to determine whether P(respond|signal) changes from tFUS were also affected by non-inhibitory behaviors. We first addressed whether tFUS within the rIFG, S1, and rIFG sham groups exerted any effects on simply responding to the Go signal by extracting mean RT from ex-gaussian distributions fit using maximum likelihood (Lacoutoure and Cousineau, 2008). Means for each subject were analyzed using a 3 x 2 mixed-design ANOVA with factors of Groups (3) and tFUS condition (2: No-tFUS, Go-tFUS trials). This analysis allowed us to assess if “going”, independent of “stopping”, was altered by potential tFUS auditory artifacts (sham rIFG group), stimulation to unrelated areas (S1 group), or whether tFUS to rIFG also influenced Go RT processes independent of a stopping context (rIFG group). We found no significant effect of tFUS condition, group, or their interaction (all p > 0.05). These results suggest that neither tFUS (rIFG and S1 groups) nor auditory factors alone (sham rIFG group) altered Go RTs independent of a Stop signal.

Improvements in inhibition from tFUS could also have emerged from reduced mean or variability in the distribution of Go reaction times occurring during stop signal trials. Two mixed-design ANOVAs were used to examine the subject-level means and variability with Group (3 levels) and tFUS (3 levels: No-tFUS, Go-tFUS, Stop-tFUS). We found no significant interactions or effects of tFUS on the mean or its variability. Together, these behavioral results indicate that only tFUS to rIFG improved response inhibition by shortening one or more processes related to the stopping speed.

### Neural responses underlying inhibition

We will now focus only on the rIFG group because this was the only group exhibiting behavioral tFUS effects. Furthermore, we only analyzed No-tFUS and Stop-tFUS conditions because Go activity was subtracted from neural data from Stop trials (see Methods). Sensor-level ERPs were examined in this group across three contrasts using cluster-based permutation t-tests: (1) successful – unsuccessful stopping contrast over each tFUS conditions, (2) successful (No-tFUS) – successful stopping (Stop-tFUS) contrast, and (3) interaction comparing successful – unsuccessful stopping between the No-tFUS and Stop-tFUS conditions. The average activity of ERP time-windows and voltage deflections of an N200 and P300 that are typically found during inhibition (Kenemans, 2015) are shown in **Figure 4A**.

**Figure 4.**
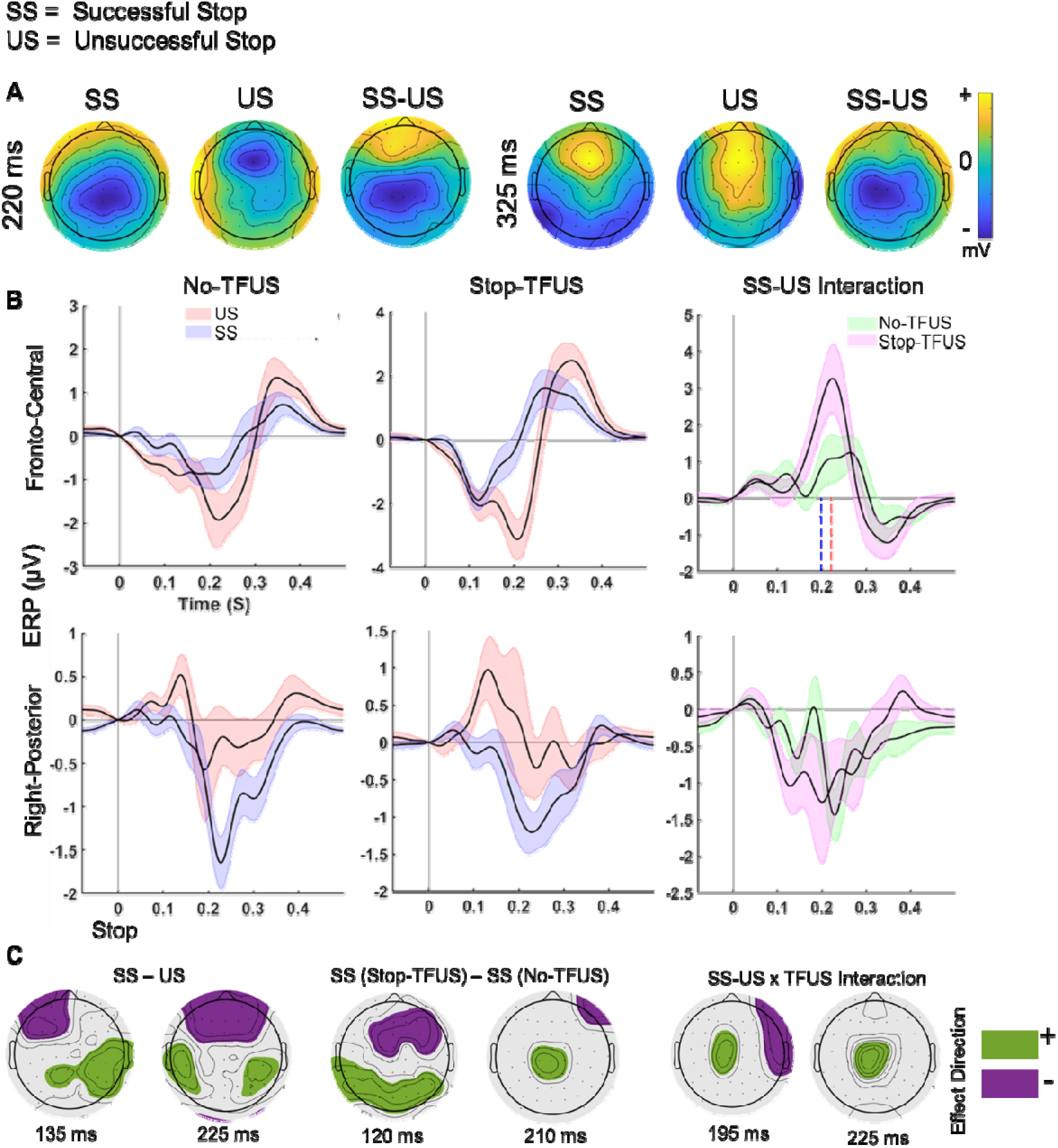
Effects of tFUS on ERPs during successful and unsuccessful response inhibition. **A.** Average scalp maps of successful (SS), unsuccessful (US), and SS-US difference maps showing ERP amplitude at time points aligned with canonical ERP effects associated with response inhibition. **B.** Average ERP plots for two a-priori chosen regions of electrodes to capture the typical N200/P300 ERP in Fronto-Central electrodes (Cz and Fcz; top row), and N100 sensory responses in Right-Posterior (CP4 and P4; bottom row). Plots show mean ERPs for SS and US trials across tFUS conditions (left and middle columns) and their interaction (right column). In the right column, the dashed vertical lines denote the median SSRT across subjects for the No-tFUS (red line) and Stop-tFUS (blue line) conditions under the interaction in Fronto-Central electrodes (top row). **C.** Permutation sensor t-tests showing the maximal effects for three comparisons: SS-US, SS trials across tFUS conditions, and interaction effect of SS-US and tFUS.

We observed an early (100-150 ms) right-posterior P100 response peaking around 135 ms that differentiated successful from unsuccessful stopping (SS and US, respectively; **Figure 4B**, left plot of **Figure 4C**). Peak location and timing are indicative of a visual P100, visible in average time-course and t-stat map (**Figure 4B-C**), with larger peak amplitudes in US trials. Contrasts of SS trials across tFUS conditions revealed the P100 was smaller on Stop-tFUS SS trials (**Figure 4C**, middle). A difference in SS trials was found for N100 responses in both frontal and right-frontal electrodes (**Figure 4C**, middle). N100s were generally larger for the Stop-tFUS trials. Neither of these ERPs exhibited an interaction of SS-US and tFUS trials. These results indicate smaller amplitudes of early sensory responses predicted successful stopping, but the lack of interaction indicates they were not directly related to inhibition.

The ERPs typically associated with indexing inhibition is the N200/P300 complex. Notably, this complex often appears in a fronto-central cluster, with the N200 peak being most prominent during failed stopping. The fronto-central ERP plot shows the N200 peaks around 200 ms but does so most clearly in US trials (**Figure 4B**, top row). Comparing SS and US trials supported this effect of larger N200 during US trials (**Figure 4C**, left). Contrasting SS trials across tFUS conditions indicated this ERP was larger for the No-tFUS condition. The interaction effect (**Figure 4C**, right) indicated the maximal difference corresponded to N200 timing, exhibiting a larger difference in the SS-US contrast during Stop-tFUS trials. N200 amplitude decreased with increasing inhibitory performance, with the smallest peak in Stop-tFUS SS trials (**Figure 4B**, top row). The fronto-central P300 amplitude also differentiated SS and US trials, with a lower amplitude for US trials. The lack of interactions for the P300 amplitude indicated it was distinct from driving the speed of inhibition (SSRT).

Our main ERP hypothesis was that P300 onset timing, rather than its peak, reflects an inhibitory process. This hypothesis stems from two considerations. The first is the N200 and P300 likely reflect a mixture of underlying components with the N200 amplitude being reflected in the P300 onset. In this case, a shift in P300 onset would drive our observed interaction contrast of the fronto-central N200 by altering the time of the N200 peak amplitude. Examination of the time course of fronto-central ERPs (**Fig. 4B**, top row) indeed indicated that reduction in N200 amplitude is likely driven by the P300 onset occurring earlier. The second consideration is that an inhibitory marker should track inhibition timing (SSRT), which has been previously reported for P300 onsets (Wessel & Aron, 2015). Therefore, we predicted that the P300 onset latency should correlate with the SSRT difference across tFUS conditions. Visually contrasting SS-US difference waveforms across tFUS conditions (**Fig. 4B**, upper right) revealed P300 onsets shifted earlier during Stop-tFUS. We found tFUS-induced changes in P300 onset significantly correlated with SSRT (r = 0.61, *p* < 0.05), thus providing support of P300 latencies tracking inhibition speed.

## Discussion

The present study builds on a rich literature that has considered response inhibition from many perspectives. By employing online tFUS to rIFG in parallel with EEG, we isolated P300 onset latencies as the primary predictor of behavioral outcomes – response inhibition performance via SSRT – during No-tFUS and Stop-tFUS trials. This result is consistent with recent work suggesting P300 onset latencies predict inhibitory outcomes and SSRTs (Huster et al., 2020; Huster, Enriquez-Geppert, et al., 2013a; Wessel & Aron, 2015), with latency modulation occurring before but close to SSRT. This close temporal proximity of neural modulation and SSRT is predicted by Stop-Signal task studies of single units in non-human primates (Hanes et al., 1998), as well as neural network (Lo et al., 2009) and accumulator models (Boucher, Palmeri, Logan & Schall, 2007; Logan, Yamaguchi, Schall & Palmeri, 2015). However, our work is the first to directly stimulate online an area in the inhibitory control network, i.e., rIFG, and show a concurrent change in P300 onset and behavior, thereby supporting their causal connection (**Fig. 4**).

Response inhibition likely involves several processes, ranging from sensory cue detection, attention, performance monitoring, and presumably explicit motor inhibition (Munakata, Herd, Chatham, Depue, Banich, & O’Reilly, 2011; Wessel & Aron, 2017; Wiecki & Frank, 2013). The observations that P300 latency modulation occurred only in the rIFG group and tFUS effects were observed only for the longer SSDs indicate that rIFG is causally involved in the response inhibition process (**Fig. 3**). The fact that the effectiveness of tFUS to rIFG in enhancing response inhibition was greater for longer SSDs further suggests that rIFG could perhaps differentiate between the Go and Stop processes. Possible explanations for not observing significant tFUS effects for the shorter SSD trials are that, even before the Go process has been initiated, the stop/inhibitory process gets initiated, or they get simultaneously initiated but the inhibitory process is prioritized based on recognition of the Stop signal according to the independent race model. This could provide less opportunity for tFUS neuromodulator effects to impact the inhibition response. In contrast, for the longer SSDs the Go process has likely begun before the Stop stimulus is displayed, requiring it to be suppressed and the inhibitory process to be initiated. Our data cannot be used to distinguish among these processes and therefore further work is needed to establish which aspect of the response inhibition dynamics is mediated by rIFG.

Consistent with other studies (Bekker et al., 2005; Kok et al., 2004), the peak amplitude of fronto-central P300 differentiated inhibitory outcomes (**Fig. 4**). However, by virtue of our experimental design, we were able to determine that that rIFG is not involved with the modulation of P300 amplitude, as indicated by the lack of difference in the P300 amplitude between the No-tFUS and Stop-tFUS conditions. This finding further supports the proposition that P300 amplitude is not a marker of response inhibition (Huster et al., 2020). It is nevertheless possible that P300 amplitude modulates non-inhibitory aspects of the response in the stop-signal task (Huster et al., 2020), e.g., outcome-monitoring and attentionally-reoriented processing that could occur after the SSRT (P300; Corbetta et al., 2008; Polich, 2007). We also found that N200 amplitude modulates with response inhibition outcomes, but this effect was limited to No-tFUS trials (**Fig. 4**). This suggests that N200 reflects non-inhibitory processes, surprise or conflict (N200; Alexander & Brown, 2011; Enriquez-Geppert et al., 2010). Lastly, the modulation of N100 in the No-tfUS condition, but not in the tFUS conditions indicates this ERP is not associated with response inhibition and may reflect sensory detection (Kenemans, 2015; Lijffijt et al., 2009).

Imaging and computational modeling studies surrounding response inhibition have shown rIFG to be a neural correlate of response inhibition (Aron 2011). However, due to the correlational nature of these techniques, the mechanisms through which rIFG is causally involved with response inhibition are still debated. Three studies have attempted to address this gap using noninvasive neuromodulation in the form of offline repetitive TMS (Chambers et al., 2006; Chambers et al., 2007; Sundby et al., 2021). All of these studies reported that neuromodulation successfully disrupted subjects’ ability to inhibit their responses, as indicated by significant lengthening of their stop signal reaction time (Chambers et al., 2006; Chambers et al., 2007; Sundby et al., 2021). However, three methodological issues should be considered in the interpretation of the TMS-induced behavioral effect: (1) the limited spatial resolution of TMS (in the order of a few mm; Passera et al., 2022) challenges its ability to isolate neuromodulation to pars opercularis/rIFG only; (2) the offline stimulation modality does not address the temporal stage(s) of response inhibition at which rIFG is involved; and (3) no significant behavioral differences were found when comparing the real and sham TMS condition, which was caused by the sham condition trials being affected by the preceding TMS trial block.

The design of the present study addresses all these limitations. First, we used a neuromodulation modality with a high spatial resolution (in the order of 1-2 mm; Tufail et al., 2010) appropriate for targeted stimulation of pars opecularis. Second, we delivered tFUS simultaneously with the Go and the Stop cue, which revealed response inhibition effects only when tFUS was delivered simultaneously with the Stop cue. Critically, the establishment of temporal control is a particularly important aspect of our design because P300 is sensitive to multiple cognitive processes, e.g., stimulus evaluation, categorization and decision making. Therefore, the contrast between Go- and Stop-tFUS behavioral effects allowed us to determine the response inhibition stage at which rIFG is involved. Third, using a spatial control outside of the inhibitory network (contralateral somatosensory cortex) allowed us to validate the specificity of tFUS behavioral effects to rIFG.

This study strengthens the evidence for tFUS as a promising neuromodulation technique by demonstrating its ability to significantly modulate P300 onset latency while tracking response inhibition. Compared to alternative neuromodulation techniques, tFUS offers higher spatial resolution, thus providing the opportunity to selectively target brain areas with high spatial specificity and assessing their causal role in the phenomenon under study (Tufail et al., 2010; Siddiqi et al., 2022). Additionally, tFUS has the ability to penetrate the skull and brain tissue to reach deep brain targets non-invasively with minimal non-target effects (Fini & Tyler, 2017). Since ultrasound is not an electromagnetic signal, tFUS does not interfere with neuroimaging techniques and can therefore be applied simultaneously with fMRI or EEG, as done in the present and previous work (e.g., Legon et al., 2014), as well as other neuromodulation techniques such as TMS (e.g., Legon et al., 2018). In recent studies, researchers are exploring its therapeutic abilities as well (Sanguinetti et al., 2020; Reznik et al., 2020).

In conclusion, competing theories have argued whether the P300 amplitude or onset latency is a marker of response inhibition and if rIFG plays an important role in modulating the P300 with respect to response inhibition. Our behavioral and ERP results favor the framework wherein P300 onset latency is a reliable marker of response inhibition success – as demonstrated by its association with stopping speed – and rIFG is causally involved in response inhibition. However, further work is needed to determine the mechanisms by which rIFG modulates the P300 onset latency. For example, rIFG may act on downstream areas linked to medial frontal P300 generation through a variety of different circuit mechanisms, such as feedforward excitation, inhibition, or extra-cellular response gain modulation. Animal model recordings from both areas, combined with tFUS, could help clarify the mechanism(s) responsible for inhibition speed changes. Another area for future investigation is determining what aspects of the response inhibition process are regulated by rIFG. We found rIFG comes into play only after the Go process has started, whereas the Stop process has to be initiated for successful inihibition. Hence, the temporality with which rIFG is involved in response control could be involved in suppressing the Go process, enhancing the Stop process or both.

Finally, given the high spatial resolution of tFUS for modulation of superficial (cortical) and deep brain networks, it has been suggested to be a valuable tool for basic neuroscience, as well as for the development of therapies in psychiatric medicine (Fini & Tyler, 2017). The use of tFUS has also been suggested to be a powerful method for mapping human brain circuitry and behavior (Tyler, Lani & Hwang, 2018). The present use of tFUS and EEG thus provides a neural and methodological framework for testing causal, underlying hypotheses and models regarding functional brain-behavioral dependencies (Siddiqi et al., 2022). The efficacy of tFUS in enhancing successful response inhibition reported here also underscores the potential use of tFUS as an intervention for behavioral disorders stemming from inhibitory response dysfunction (e.g., impulsivity, ADHD, and substance abuse disorder; Bari and Robbins, 2013). Our data and observations indicate that future studies combining tFUS with behavioral assays and neurophysiology can uncover additional targets and approaches for studying and treating a variety of neuropsychiatric conditions and thought disorders caused by aberrant activity in cortical and deep-brain networks.

## Data availability

The human behavioral and EEG ERP datasets for reproducing figures are publicly available. The datasets are publicly available on Dryad: shorturl.at/BOWX6 Dryad Digital Repository, doi:10.5061/dryad.sj3tx968j

## Code availability

Behavioral and ERP Analysis scripts for reproducing figures are available at: https://github.com/justfineIU/DCM_MODS.

Other codes for analysis are available from the corresponding author upon request.

